# Flagellin triggers mesophyll dehydration: An early PTI defense against bacterial establishment in intercellular spaces

**DOI:** 10.1101/2020.12.31.424953

**Authors:** Ahan Dalal, Ziv Attia, Menachem Moshelion

## Abstract

Plants have evolved various mechanisms to defend themselves against pathogens. Many pathogens induce the formation of water-soaked lesions during early infection under conditions of high atmospheric humidity. These water-soaked spots are caused by the disruption of the plasma membrane or cell wall integrity due to various activities of effector proteins during infection. We hypothesized that bacterial PAMP-flagellin plays a role in modulating the cell-membrane permeability that controls the availability of water in the apoplast, to prevent bacterial establishment on the cell wall during the early stages of the PAMP-triggered immunity (PTI) response. Our results revealed that the conductivity of hydraulic pathways in the leaf was reduced in response to flagellin22 (flg22). The cellular osmotic water permeability (P_f_) of both mesophyll cells and bundle-sheath cells was dramatically reduced in response to flg22 treatment. Moreover, the whole-leaf hydraulic conductance (K_leaf_) was also reduced in response to flg22 treatment. The fact that the P_f_ of mesophyll cells of an aquaporin (AQP) mutant was not affected by the flg22 treatment suggests the involvement of AQP channels in the flg22-induced P_f_ reduction signal transduction pathway. We conclude that the binding of flagellin to their receptors elicits signals to close AQPs, consequently reducing the water content of the cell wall and intercellular spaces and leading to a more negative water potential. This serves as an early PTI response to pathogen attack, which, in turn, might decrease the rate of bacterial growth and establishment in the apoplast.

**Significance statement:** We report that the membrane osmotic water permeability of both leaf mesophyll and vascular bundle-sheath cells is reduced in response to treatment with flagellin22. We suggest that this mechanism for cell dehydration may serve as an apoplastic defense response, to limit the chances of bacterial pathogens becoming established on the walls of leaf mesophyll cells.

## INTRODUCTION

Plants are continuously exposed to various kinds of pathogens. The physical barriers of the epidermal cuticle and cell wall serve as a first line of defense, which is also biochemical in nature, in that those organs constitutively produce antimicrobial compounds. Unlike fungal pathogens, bacteria can only enter plants through natural openings or wounds and cannot directly penetrate the healthy plant epidermis (reviewed by [1]). Even before the pathogen manages to enter the plant, the plant’s immune system is activated. Pathogens are recognized by their conserved signature components, known as pathogen-associated molecular patterns (PAMPs). PAMPs are recognized by the plant’s immune receptors, with extracellular domains known as pathogen or pattern-recognition receptors (PRRs), and this recognition elicits PAMP-triggered immunity or pathogen-triggered immunity (PTI).

### Stomata and apoplast: Two layers of defense

The movement of pathogens into the plant and their propagation in the plant are restricted by two major immunity mechanisms: stomatal-closure defense and apoplastic defense [2]. Stomatal defense is the first line of immune response and is known to be a part of PTI, which involves various channel activities that control stomatal closure [2□11]. Once the pathogen manages to pass through the stomatal barrier and enter the intercellular spaces of the mesophyll, it encounters an apoplastic defense barrier that restricts its propagation [2, 12, 13]. The apoplast is the space in which PAMPs are initially recognized by PRRs [7, 14]. Plant genomes contain a great number of potential PRRs, but only a few receptor-ligand pairs have been identified to date [4, 15]. This apoplastic defense system involves both PTI and another layer of immune response called effector-triggered immunity (ETI), which occurs when pathogens are perceived via the proteins, called effectors, which they secrete in the apoplast and inside host cells. Plants recognize these effectors via immune receptors that are encoded by disease-resistance genes (R genes) and this triggers an ETI response ([2, 7], reviewed by [16]).

### Cellular perception of flg22

Flagellin peptide (flg22) is a bacterial PAMP that has been widely studied in the context of PTI defense mechanisms. This peptide is a fragment of bacterial flagellin that directly binds to a transmembrane receptor kinase, FLAGELLIN SENSING2 (FLS2), which is located on the plasma membrane. In *Arabidopsis thaliana*, multiple other kinases work together with FLS2 in the defense-signaling process (reviewed by [17]). Flg22 binding induces FLS2 heteromerization with BRASSINOSTEROID INSENSITIVE 1–associated kinase 1 (BAK1). Besides directly interacting with FLS2, BAK1 also acts as a coreceptor, recognizing the C-terminus of the FLS2-bound flg22 [18]. Some Arabidopsis FLS2 is present in FLS2-FLS2 complexes before and after exposure to flg22, but flg22-binding capability is not required for FLS2-FLS2 association [17]. FLS2 binds flg22 independently of its association with BAK1. However, BAK1 is a positive regulator of PAMP signaling in Arabidopsis (19). Moreover, in that species, *BAK1* is expressed at detectable levels in both mesophyll cells (MCs) and the vascular bundle-sheath cells (BSCs) of the leaf vascular system, as seen in a transcriptome analysis [20].

### Pathogens trigger aqueous apoplast

The apoplastic defense system is regulated through the tight control of various components, namely, water, nutrients, pH, reactive oxygen species (ROS) and secreted proteases and antimicrobial compounds (reviewed by [1, 21]). The availability of water in the apoplast determines the success of pathogen propagation and infection. Many pathogens have been known to induce the formation of water-soaked lesions during the early infection phase under conditions of high atmospheric humidity ([22, 23], reviewed by [24]). These transient spots are the regions in which bacteria aggressively proliferate and the spots disappear before the appearance of late disease symptoms [24, 25]. These water-soaked spots are formed due to aqueous apoplast caused by the disruption of the plasma membrane or the integrity of the cell wall integrity, as a result of various activities of the effector proteins during infection (reviewed by [24]).

Many studies have examined early defense signaling in apoplast. One of the early signaling pathways in the mesophyll cells of Arabidopsis is the flg22-mediated, BAK1-dependent, calcium-associated plasma membrane anion channel opening pathway [26]. Findings suggest that activation of ETI can block the water-soaking process, possibly as an integral part of the plant defenses against bacterial pathogenesis (reviewed by [24]). However, the effect of PAMP in the inhibition of apoplast “over-wetting” has not been studied.

In this study, we hypothesized that flagellin22 plays an important role in modulating cell-membrane permeability, thereby regulating the availability of water in the apoplast, to prevent bacterial establishment on the cell wall during the early stages of a PTI response.

## RESULTS

The permeability of the cell membrane (which is regulated by various factors) determines the transient water status of the cell wall and its surrounding apoplast. We measured the effect of flg22 on the permeability to water (P_f_) of MC protoplasts from WT plants, *bak1-4* mutants and mutant plants in which the mutation was complemented by the *BAK1* gene driven by its native promoter (BAK1pro::BAK1/*bak1-4*). When mesophyll protoplasts were subjected to hypotonic challenge, we observed slower swelling of the flg22-treated WT cells (Fig. 1A), due to the significantly lower water permeability of their membranes (Fig. 1B). The *bak1-4* mutant did not exhibit any reduction in its swelling in response to flg22 treatment (Fig. 1C), reflecting the fact that the flg22 treatment did not impair its membrane water permeability (Fig. 1D). The complemented-mutant cells (BAK1pro::BAK1/*bak1-4*) revealed a gain of functional behavior in their swelling in response to hypotonic challenge (Fig. 1E), as well as a significant reduction in the permeability of their cell membranes in response to flg22 treatment (Fig. 1F).

**Figure 1.**
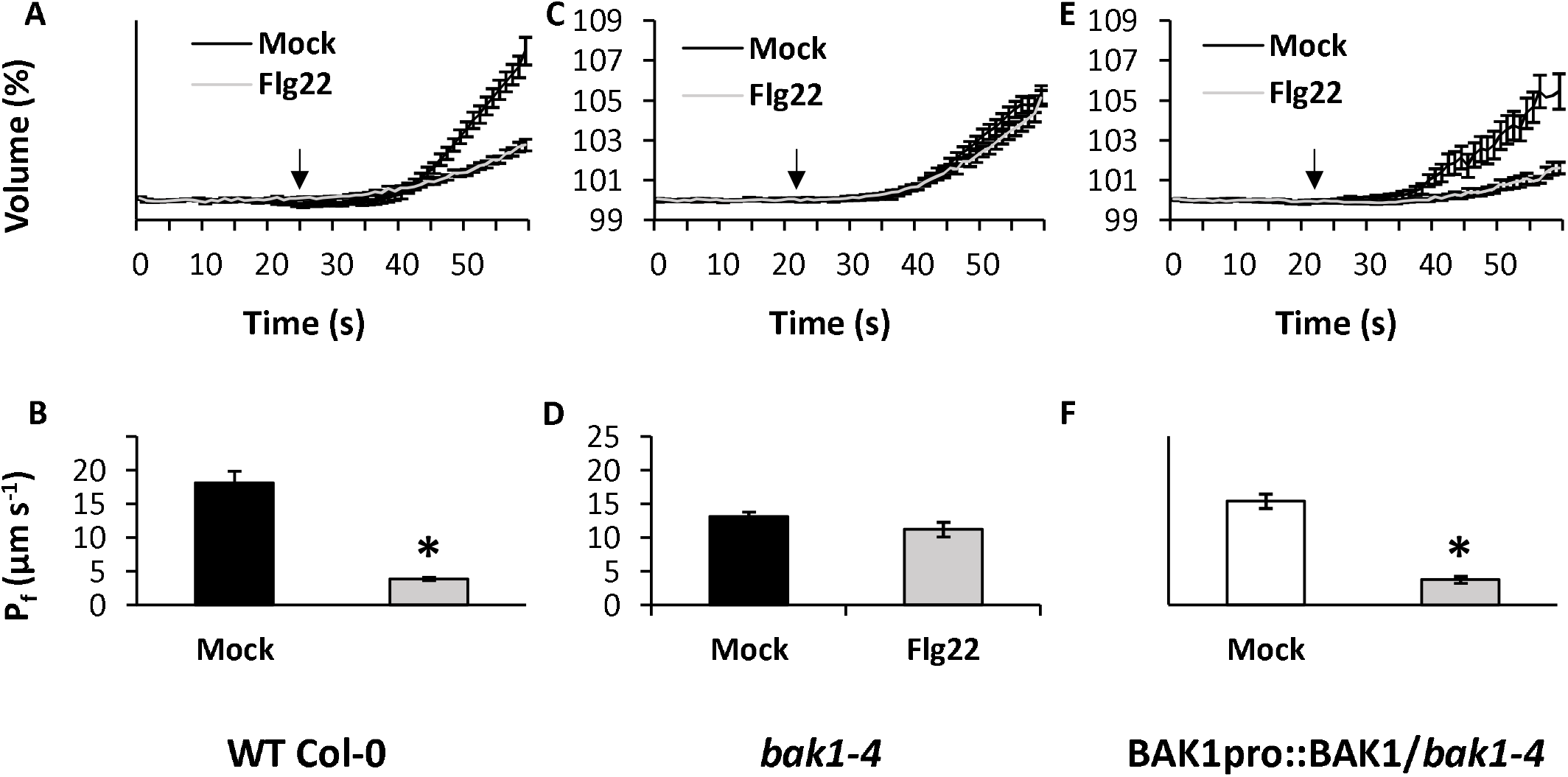
The effect of flg22 on the cellular osmotic water permeability (P_f_) of the protoplasts from Arabidopsis mesophyll cells. **(A, C, E)** The increase in the volume of the different protoplasts: wild type (WT Col-0), *bak1-4* mutants and the mutants complemented with *BAK1* gene driven by its native promoter (BAK1pro::BAK1/*bak1-4*) over 60 s during the hypotonic challenge. The arrows in each graph indicate the starting time. **(B, D, F)** P_f_ of the WT Col 0, *bak1-4* and BAK1pro::BAK1/*bak1-4* protoplasts. Data are shown as means ± SE. Asterisks represent significant differences between the treatments (*t*□test, *P* < 0.05).

Water channels known as aquaporins (AQPs) are known to regulate membrane water permeability (P_f_) [27]. Constitutive silencing of the PIP1 family of AQPs was shown to reduce the PIP1 (entire family) and transcript levels of several PIP2 genes, as well as to suppress the P_f_ of bundle-sheath and mesophyll cells [27]. In this study, we used the same PIP1-silenced line (35S:mirPIP1-8) that Sade et al. [27] used to investigate whether PIP1 plays a role in flg22-mediated reduction of mesophyll P_f_. In our study, hypotonically challenged mesophyll protoplasts did not show any reduction in swelling in response to flg22 treatment (Fig. 2A), reflecting the fact that the permeability of their cell membranes to water was not reduced by the flg22 treatment (Fig. 2B).

**Figure 2.**
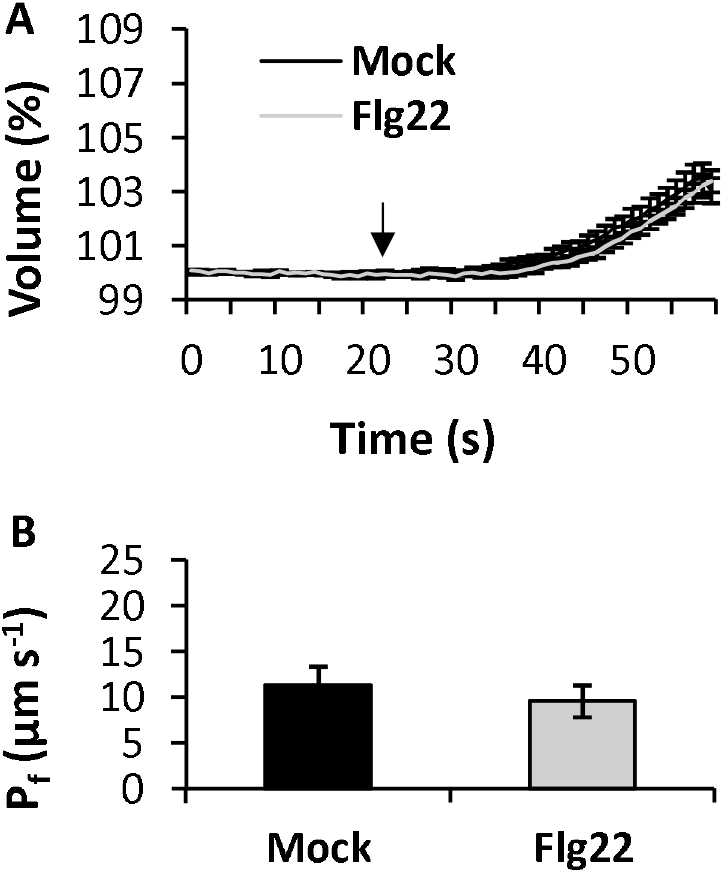
The effect of flg22 on the cellular osmotic water permeability (P_f_) of the protoplasts from mesophyll cells of PIP1-silenced (35S:mirPIP1) Arabidopsis. **(A)** The increase in the volume of the protoplasts over 60 s during the hypotonic challenge. The arrow points to the starting point. **(B)** P_f_ of the protoplasts. Data are shown as means ± SE. Asterisks represent significant differences between the treatments (Student’s *t*□test, *P* < 0.05).

Water enters the leaf from the xylem through the BSCs and then moves into the intercellular space, in a pattern known as radial water influx (Kleaf). BSCs form a layer of living parenchyma cells that tightly enwraps the dead xylem. To investigate the effect of flg22 on the P_f_ of BSCs, we used green fluorescent protein (GFP)-labelled BSCs from transgenic SCR::GFP Arabidopsis plants. Hypotonic-challenged bundle-sheath protoplasts revealed slower swelling following flg22 treatment (Fig. 3A), due to the lower water permeability of their membranes (Fig. 3B). The P_f_ of BSCs regulates K_leaf_ [28]. To measure K_leaf_, we allowed the leaves to take in the treatment solution through their petioles (xylem-fed). Before measuring K_leaf_, we checked whether the leaves respond to flg22 treatment by confirming (a) *AtWRKY33* expression in WT and (b) GUS expression in MYB51pro::GUS transgenic Arabidopsis leaves using the same petiole-feeding experimental set-up. Quantitative PCR (qPCR) results showed that the expression level of *AtWRKY33* was significantly higher in the treated plants, as compared to the untreated mock (Supplementary Fig. 1A). The response assay revealed an increase in GUS staining throughout the leaf upon treatment, implying that the flg22 response is transmitted beyond the vascular tissues (Supplementary Fig. 1B). Earlier studies have shown that *AtWRKY33* and *AtMYB51* expression is induced in wild-type (WT) Arabidopsis in response to treatment with flg22 [29, 30]. Flg22 has also been shown to elicit the expression of the β-glucuronidase (GUS) reporter gene driven by MYB51 promoter (MYB51pro::GUS) in the root of transgenic Arabidopsis [30]. We showed that flg22-treated leaves exhibited significantly reduced K_leaf_ (by 56%), as compared to the non-treated leaves. In contrast, the *bak1-4* mutants did not seem to respond to the flg22 treatment and did not show any change in their K_leaf_ (Fig. 3C). The higher calculated K_leaf_ under mock conditions (without flg22) was due to a much greater (less negative) leaf water potential, Ψ_leaf_ (Materials and Methods; Fig. 3E). This invites the interpretation that flg22 slowed the movement of water into the leaf, reducing the radial water influx which, in turn, limited transpiration (Fig. 3D).

**Figure 3.**
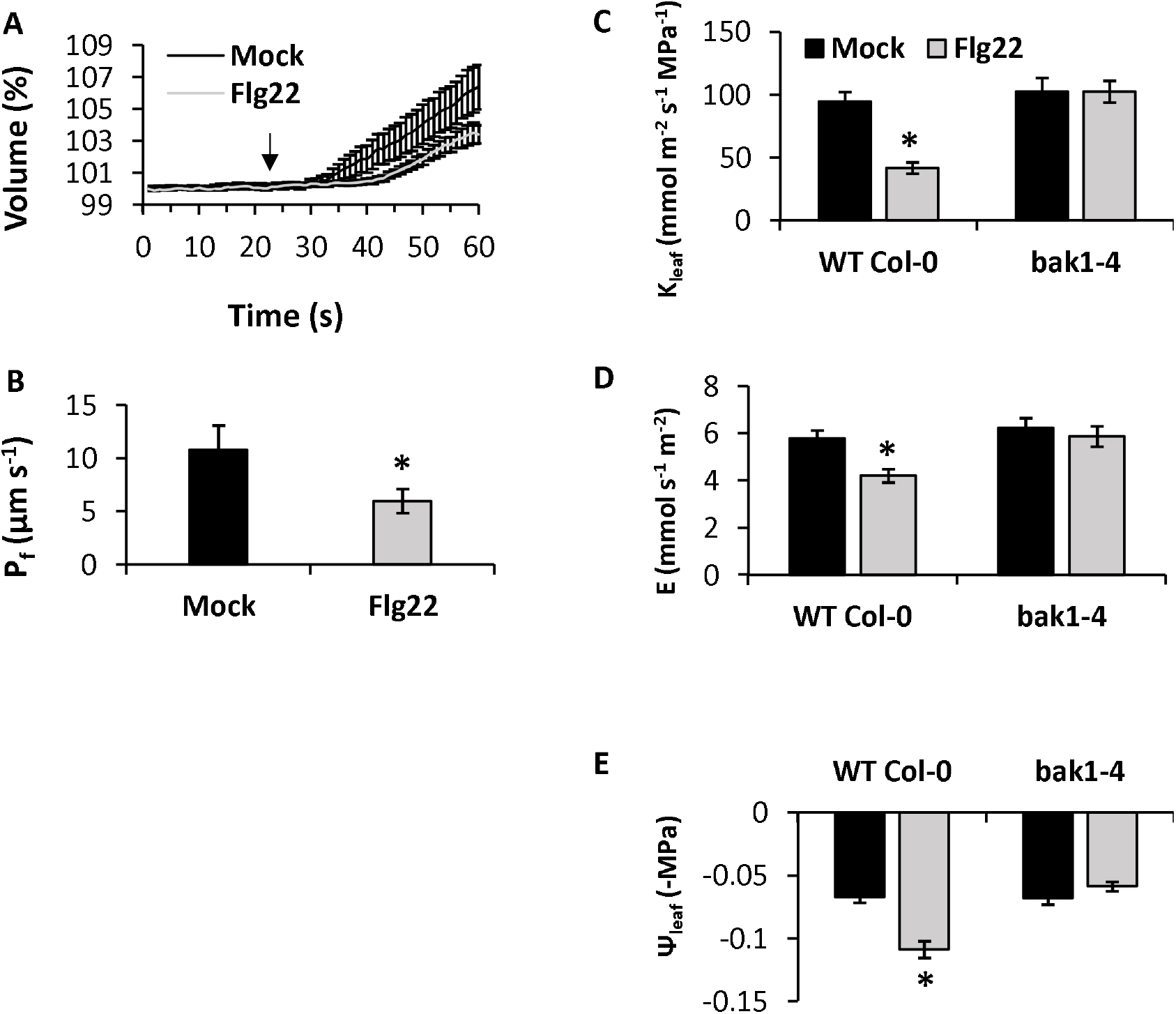
The effect of flg22 on the cellular osmotic water permeability (P_f_) of the bundle-sheath protoplasts from **(A, B)** wild-type plants and the relative change in **(C)** leaf hydraulic conductance, K_leaf_, **(D)** transpiration rate, E, and **(E)** leaf water potential, Ψleaf, in wild-type (WT Col-0) and *bak1-4* mutant protoplasts. Data are shown as means ± SE. Asterisks represent significant differences between treatments within the same genotype (Student’s *t*□test, *P* < 0.05). Data are from at least three independent experiment days.

## DISCUSSION

Many bacterial pathogens induce apoplast over-wetting to thrive and propagate in the apoplastic space during the early infection phase [24]. In this study, we tested whether there is an early PTI response that regulates intercellular moisture. Intercellular moisture is an important factor that determines bacterial growth during hypersensitive response [31], which then inhibits bacterial establishment and proliferation on the cell wall. The water stress experienced by bacteria (with flagella) in the mesophyllic apoplast may be part of an early flagellin-induced PTI response that prevents bacterial establishment on the cell wall.

### Flagellin regulates the hydration of the mesophyllic apoplast

Water enters the leaf from the xylem through BSCs, which function as a selective barrier that allows the intake of water and solutes [32]. Under optimal conditions, the MC wall is wet due to both symplastic (via the mesophyll tissue) and apoplastic (the continuum of radial flow of water from the xylem vein) water flow. Our results revealed that these hydraulic pathways in the leaf were reduced in response to treatment with flagellin. The cellular osmotic water permeability (P_f_) of the mesophyll cells and the bundle-sheath cells was dramatically reduced in response to flg22 treatment (by 78% and 45%, Fig. 1B and Fig. 3B, respectively). Moreover, the whole-leaf hydraulic conductance (Kleaf) was reduced in response to flagellin applied through the xylem (Fig. 3C). This reduction occurred despite the fact that we performed the flg22 treatment and gas-exchange measurements in a high-humidity environment (low vapor pressure deficit ~ 1.35□1.45 KPa, as in Attia et al. [33]), in order to mimic the optimal conditions for bacterial infection conditions [23□25],

Lower P_f_ of the BSC is known to restrict the movement of water into the leaf, which reduces the leaf water potential (reviewed by [34]), as confirmed by our results (Fig. 3E). Interestingly, lower water potentials have been strongly correlated with smaller bacterial population sizes *in planta* [31]. This supports our hypothesis that the induction of a more negative leaf water potential might be an effective tool in the plant’s initial response to bacterial pathogens. Similarly, chitin, which is a fungal PAMP, has also been shown to reduce P_f_ in the mesophyll and bundle-sheath cells of Arabidopsis [33]. In addition, Li et al. [35] reported an observation with another bacterial PAMP named harpin, which promotes cell hydraulic conductivity in leaves and roots of WT Arabidopsis. Thus, we conclude that this initial apoplastic dehydration phase may slow or restrict bacterial cell division, which could provide the plant with more time to activate PTI defense-signaling cascades.

### Cellular mechanism controlling the early P_f_-regulated PTI response

BAK1 is the co-receptor for flagellin. Therefore, we checked whether there were any flg22-induced P_f_ changes in *bak1-4* mutants or the complemented mutants (BAK1pro::BAK1/*bak1-4*). We observed a loss-of-function (no change in P_f_) with flg22 treatment in *bak1-4* mutants (Fig. 1D), which was similar to that observed previously in the guard-cell protoplasts of *bak1-5* mutants [36]. However, the lost function was regained in complemented BAK1pro::BAK1/*bak1-4* plants, which exhibited a significant (75%) reduction in P_f_ following flg22 treatment (Fig. 1F). Similarly, in earlier studies in which the leaves of WT plants, *bak1-4* mutants and the complemented mutants (BAK1pro::BAK1*/bak1-4*) were checked for total ROS burst (indicative of a response to flg22), the response to flg22 was found to be impaired in *bak1-4* mutants and recovered in BAK1pro::BAK1/*bak1-4* plants [19, 37].

P_f_ is regulated by AQPs [27]. Many studies have found that AQPs play roles in infection and immunity in plants ([34, 38□41]; reviewed by [42]). PIP AQPs have extracellular regions that are exposed to the outside of the plasma membrane [39] and can perceive abiotic and biotic stress signals (reviewed by [42]). Moreover, harpin, a bacterial PAMP, induces PIP function through direct interaction during defense or infection in plants [35, 38]. The relative amounts of water moving through the three different transmembrane pathways (i.e., AQPs, the symplast and the apoplast) may be altered by AQPs [43]. We observed no change in the P_f_ of 35S:mirPIP1-8 MCs in response to flg22 treatment (Fig. 2B), which suggests that AQPs are involved in the flagellin-induced reduction of MC membrane permeability. These results support our hypothesis that when MCs encounter flagellin, an early PTI response is triggered in which flagellin modulates the PIPs and restricts the permeability of the cell membrane, thereby inducing apoplast dehydration. Nevertheless, wetting of the cell wall may be also related to the apoplastic flow of fluid/water coming from the vascular system. This radial flow known as K_lea_f is reportedly regulated by the vascular bundle sheath [27, 28].

Rodrigues et al. [36] showed that flg22 activates AtPIP2;1 to increase the P_f_ of guard-cell protoplasts, resulting in turgor loss and stomatal closure. Stomatal closure is considered to be the first line of defense against bacterial pathogens. We propose that this flg22-induced PIPs in the bundle sheath, mesophyll, and guard-cell protoplasts act in series during early infection to prevent bacterial propagation.

## CONCLUSION

Here, we provide evidence that suggests the possibility of a PTI-related response that induces leaf dehydration, which might create a less favorable microenvironment for bacterial pathogens, which could slow bacterial establishment in the leaf (Fig. 4). Once the bacteria manage to enter the wet intercellular spaces, their flagellin is recognized by the receptors on the plasma membranes of mesophyll and vascular bundle-sheath cells. The binding of flagellin to these receptors might elicit signals to close aquaporins and reduce the osmotic water permeability of the MC and BSC membranes, as well as Kleaf. This could reduce the water content of the cell wall and intercellular spaces. Continuous transpiration reduces the amount of water in the leaf apoplast even further, which may inhibit the establishment of bacteria and help the plant to buy more time to initiate other defense pathways. Future research might open up new avenues in understanding how plant water relations shape the PTI response to pathogens at early stages of infection.

**Figure 4.**
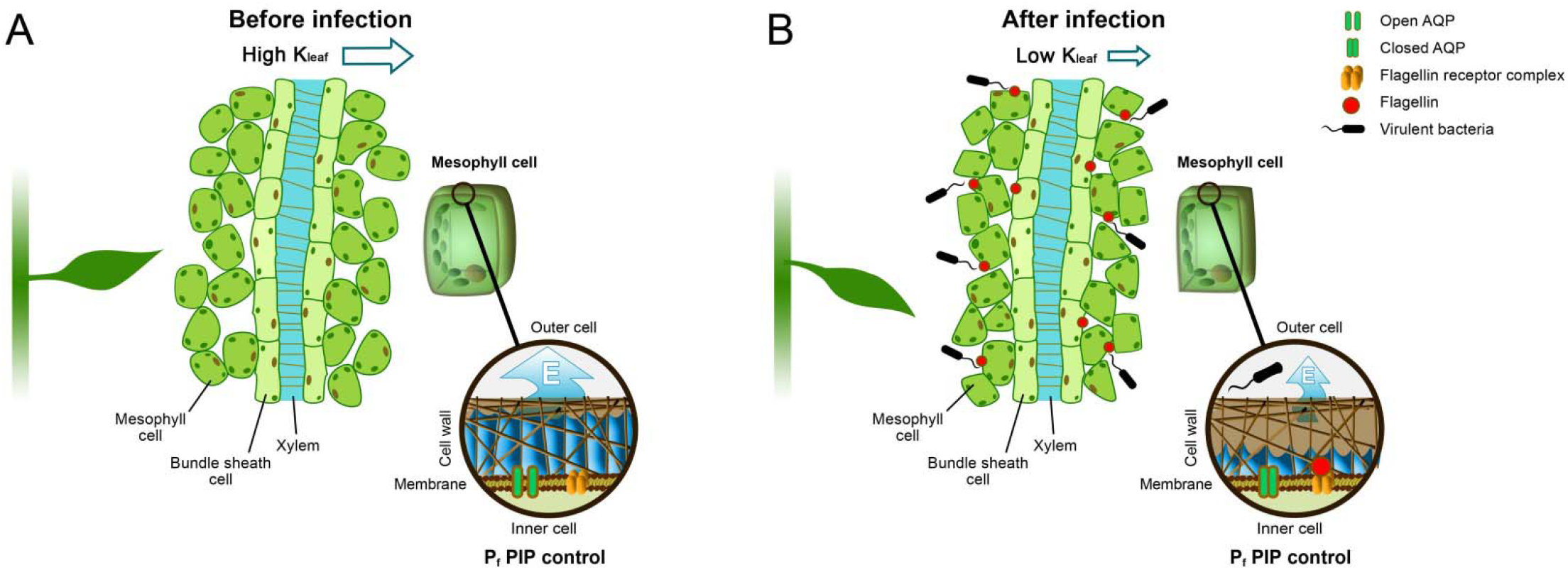
Schematic diagram of early defense response of flagellin-elicited apoplastic dehydration of mesophyll cells. In this hypothetical model, we present a primary PTI response triggered in the early stages of pathogen entry, which prevents the cell wall from becoming too wet, in order to inhibit bacterial establishment. **(A)** Under well-irrigated conditions, relatively high leaf-hydraulic conductance (movement of water from the xylem into the mesophyll through the bundle-sheath layer) maintains a relatively high (close to zero) water potential, keeping intercellular spaces wet, which enables high levels of transpiration. **(B)** Once the bacteria cross the stomatal barrier, they enter the wet intercellular space, where they can colonize and thrive. Flagellin released from bacteria in the intercellular apoplastic spaces is recognized by receptors on the plasma membranes of mesophyll and vascular bundle-sheath cells. The binding of flagellin to these receptors elicits signals to close the aquaporins. Transpiration or water loss from the surface of the leaf while water flow from xylem is restricted will contribute to additional apoplastic dehydration. This gradually reduces the amount of water inside the cell walls and in intercellular spaces [43] and generates a more negative water potential within the microcapillary structure of the mesophyll cell walls [46]. Lower water potentials, due to apoplastic dehydration, can slow down bacterial cell division, which may be a feasible plant defense strategy as it provides the plant with more time to activate other signaling cascades of the PTI defense response.

## MATERIALS AND METHODS

### Plant materials and growing conditions

This work was conducted using *Arabidopsis thaliana* plants, specifically *A. thaliana* Col-0 (WT), the T-DNA insertion mutant line *bak1-4* (SALK_116202C), the complemented mutants BAK1pro::BAK1/*bak1-4* and MYB51pro::GUS (N68983; MYB51 promoter-driven GUS; [33]), the PIP1-silenced line 35S:mirPIP1-8 [27] and SCR::GFP (SCARECROW (SCR) promoter-driven GFP; [33]) plants. Seeds of *bak1-4* and BAK1pro::BAK1/*bak1-4* plants were a generous gift from Prof. Cyril Zipfel, University of Zurich, Switzerland.

All plants were germinated and grown in soil (Klasmann686 Klasmann-Deilmann, Germany) containing slow-release fertilizer (4 g/L; Osmocote^®^ 6M). The plants were grown under short-day conditions (8 h light/16 h dark) at 22°C and 70% relative humidity. LED light strips [EnerLED 24 V-5630, 24 W/m, 3000 K (50%)/6000 K (50%)] were used for illumination with an intensity of 150 200 μmol m^-2^ sec^-1^ when measured at the height of the plants’ rosettes. During the day, the automated changes in illumination, humidity and vapor pressure deficit resembled natural conditions. The plants were irrigated twice a week [44].

### Flg22: Preparation and treatment

Flagelin22 (flg22) (GenScript) stock solution (1 mM) was prepared in sterile double-distilled water. A final concentration of 1 μM was used for all of the experiments. For the measurement of osmotic water permeability (*P*_f_), protoplasts isolated in isotonic solution (600 mosmol) were treated with flg22 (mock left untreated) for 2–4 h in round-bottom, 2-ml centrifuge tubes prior to the measurements.

For measurements of leaf hydraulic conductance (K_leaf_), transpiration (E) and water potential (Ψ_leaf_), the flg22 was applied using the petiole-feeding set up described in Attia et al. [33], with a few modifications. Briefly, fully expanded leaves of similar size (approximately 6 cm^2^) with similar vascular areas and no noticeable injuries or anomalies were excised from 6–7 week□old plants at the petiole base with a sharp blade in the dark (pre-dawn). The harvested leaves were immediately immersed (petiole dipped) in 0.6 ml centrifuge tubes containing artificial xylem solution (AXS, for composition see [33]) with or without flg22. The tubes containing the leaves were transferred to a transparent box (3.3□L, 20 × 20 × 8.25 cm) with moist tissue paper placed on the bottom of the box to maintain a humidity level of ~90%. The closed boxes (with transparent lids) were incubated under continuous light in the same growth chamber in which the plants were grown previously for 2–4 h before the measurements were taken.

### RNA isolation and qPCR

To analyze the transcript expression of the flg22□induced marker gene *AtWRKY33* in WT plants, flg22 treatment was applied via the petiole-feeding set up described in the Flg22 Preparation and Treatment section above. Total RNA was isolated from *A. thaliana* leaves and subsequent cDNA synthesis, qPCR and relative gene expression analyses were performed using the same kits, equipment and protocols described in Attia et al. [33]. The primers that we used are listed in Supplementary Table 1.

### GUS assay

To analyze the flg22 response, we used MYB51pro::GUS Arabidopsis plants. Flg22 treatment was applied as described in the Flg22 Preparation and Treatment section above. The GUS assay was performed using the same kits, equipment and protocols described in Attia et al. [33].

### Protoplast isolation and measurement of osmotic water permeability (P_f_)

Protoplast isolation and P_f_ determination were performed as described by Shatil-Cohen et al. [45] and Attia et al. [33]. Briefly, protoplasts were isolated from 6-to 7-week-old BSCs and the protoplasts from SCR::GFP plants was screened for GFP activity. P_f_ was measured by exposing single protoplasts of both MCs and BSCs to hypo□osmotic challenge (transfer from a 600□mosmol isotonic bath solution to a 500□mosmol hypotonic solution). P_f_ was calculated toward the end of the hypo□osmotic challenge.

### Determination of E, Ψ_leaf_ and K_leaf_

For the measurement of E, Ψ_leaf_ and K_leaf_, we followed a protocol similar to the one described by Attia et al. [33]. Briefly, the treatment boxes were opened for 3 min and the leaves were placed in a LI□COR 6800 gas□exchange system (LI□COR, https://www.licor.com/). The illumination settings in the cuvette were set to the same conditions as those found in the growth room. After measuring E, the water potential Ψ_leaf_) was determined immediately using a pressure chamber (ARIMAD-3000; MRC Israel) and K_leaf_ was calculated as follows: K_leaf_ = -E / (**Ψ**_leaf_ - **Ψ**_AXS_) ≈ -E / **Ψ**_leaf_. All measurements were taken between 10:00 and 13:00.

### Statistical analysis

The statistical analysis was performed using JMP 15.0 software. The means were compared using Student’s *t*□test and were considered significantly different at *P* < 0.05.

## ACKNOWLEDGEMENTS

This research was supported by the Israel Science Foundations (ISF grant no. 1043/20 to MM). AD and MM thankfully acknowledge the United States–Israel Binational Science Foundation (BSF, grant no. 2015100) for the postdoctoral scholarship awarded to AD.

## AUTHOR CONTRIBUTIONS

MM conceived the original research plan. AD and ZA performed the research and analyzed the data. AD, ZA and MM wrote the article. All authors were involved in reviewing and editing the manuscript.

