## Supplementary Figure for "Flagellin triggers mesophyll dehydration: An early PTI defense against bacterial establishment in intercellular spaces"

### Slide 1
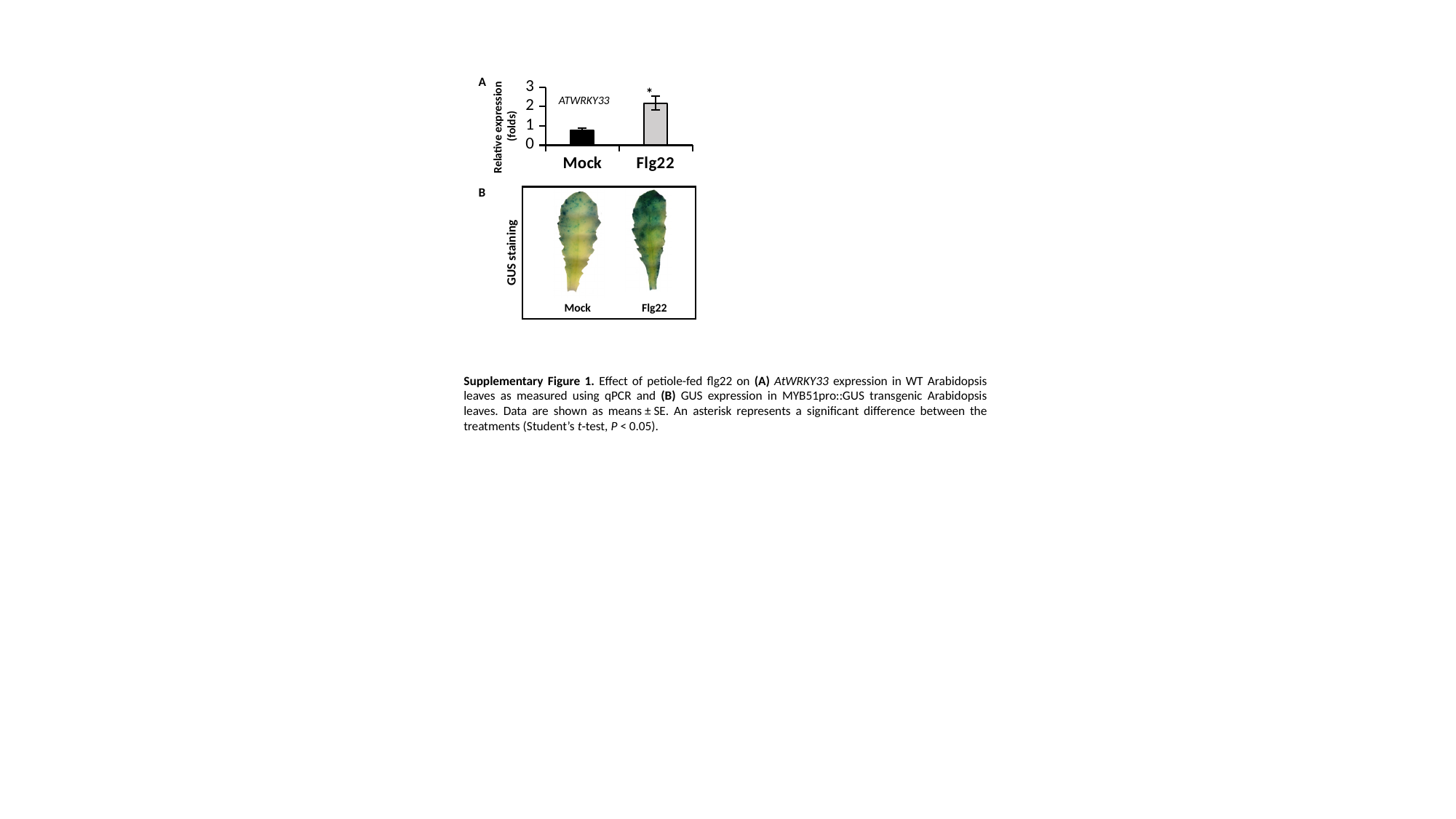

A
#### Chart
| Category | |
|---|---|
| Mock | 0.783807203 |
| Flg22 | 2.1823503933 |*
ATWRKY33
Relative expression
(folds)
B
GUS staining
Mock
Flg22
Supplementary Figure 1. Effect of petiole-fed flg22 on (A) AtWRKY33 expression in WT Arabidopsis leaves as measured using qPCR and (B) GUS expression in MYB51pro::GUS transgenic Arabidopsis leaves. Data are shown as means ± SE. An asterisk represents a significant difference between the treatments (Student’s t‐test, P < 0.05).
