## Supplementary Table for "Flagellin triggers mesophyll dehydration: An early PTI defense against bacterial establishment in intercellular spaces"

**Supplementary Table 1. Primers used for qPCR**

| **Gene** | **Primer name** | **Primer sequence** |
| --- | --- | --- |
| *WRKY33* | WRKY33-F:  WRKY33-R: | 5’ TACGAAGGGAAACACAACCA 3’  5’ AAGGCCCGGTATTAGTGTTG 3’ |
